# “Role of a Pdlim5:PalmD complex in directing dendrite morphology”

**DOI:** 10.1101/2023.08.22.553334

**Authors:** Yogesh Srivastava, Maxsam Donta, Lydia L. Mireles, Adriana Paulucci-Holthauzen, M. Neal Waxham, Pierre D. McCrea

**Author notes:** Co-corresponding authors: Pierre D McCrea PhD, Yogesh Srivastava PhD, M Neal Waxham PhD.

## Abstract

Neuronal connectivity is regulated during normal brain development with the arrangement of spines and synapses being dependent on the morphology of dendrites. Further, in multiple neurodevelopmental and aging disorders, disruptions of dendrite formation or shaping is associated with atypical neuronal connectivity. We showed previously that Pdlim5 binds delta-catenin and promotes dendrite branching (Baumert et al., *J Cell Biol* 2020). We report here that Pdlim5 interacts with PalmD, a protein previously suggested by others to interact with the cytoskeleton (*e.g*., *via* adducin/spectrin) and to regulate membrane shaping. Functionally, the knockdown of PalmD or Pdlim5 in rat primary hippocampal neurons dramatically reduces branching and conversely, PalmD exogenous expression promotes dendrite branching as does Pdlim5. Further, we show that each proteins’ effects are dependent on the presence of the other. In summary, using primary rat hippocampal neurons we reveal the contributions of a novel Pdlim5:PalmD protein complex, composed of functionally inter-dependent components responsible for shaping neuronal dendrites.

## INTRODUCTION

The shaping of dendrites enables the formation of elaborate dendritic arbors required in constructing neuronal networks ^1,2^. Further, in multiple pathological contexts, the gross morphology of dendrites is altered in ways that degrade nervous system functions. Such losses come about since neuronal activity, signaling, and plasticity each require organized synaptic connections that are dependent upon the morphology of dendrites.

Constituting a gap in knowledge, less is known about the mechanisms regulating dendritic branching or lengthening relative to the wealth of studies addressing axon trajectories ^3,4^, or the formation of dendritic spines or pre- or postsynaptic densities ^5,6^. In the context of neuronal dendritic branching, we recently reported that Pdlim5 (as well as Magi1) interact with delta-catenin in a phosphorylation-dependent manner and regulate dendritic branching ^7^. The mouse knockout of Pdlim5 is embryonic lethal ^8^ and Pdlim5’s structure and presence is largely conserved across vertebrate animals (Supplemental Figure 1A) (e.g., ∼89% identity between human and rat). Dysregulation of Pdlim5 is proposed to factor into atypical dendritic tree morphology and synaptic connectivity in human neurodevelopmental diseases including schizophrenia, depression, and bipolar disorder ^8-13^ consistent with our in vitro studies identifying a regulatory role for Pdlim5 in dendrite morphology ^7^.

Pdlim5 is composed of an N-terminal PDZ-domain (*P*SD-95, *D*lg, and *Z*O-1), central DUF-domain (“Domain of Unknown Function”), and a triad of C-terminal LIM-repeats (LIM domain) (Figure 1B) ^14,15^. Pdlim5 is present in dendrites and other neuronal structures, with its subcellular distribution varying somewhat across developmental stages ^7,16^. Pdlim5 is reported to associate with alpha-actinin and PKCepsilon and to be required in phorbol ester induced dendritic growth-cone collapse, restricting dendrite outgrowth in that context ^15,16^. By having an inhibitory effect upon Rac to displace Arp2/3 from the leading edge, Pdlim5 also limits AMPK-stimulated cell migration in C2C12 cells via restriction of lamellipodia formation ^17^. In excitatory pyramidal hippocampal neurons, Pdlim5 inhibits formation of dendritic spines (via the inhibition of SPAR) ^14^. In relation to gene activity, Pdlim5 affects neuron differentiation through cytoplasmic sequestration of the Id2 transcriptional regulator ^18^. In summary, Pdlim5 exhibits context-dependent positive and negative roles in dendrites, with Pdlim5 expression increasing dendritic arbor complexity as we have reported ^7^.

**Figure 1.**
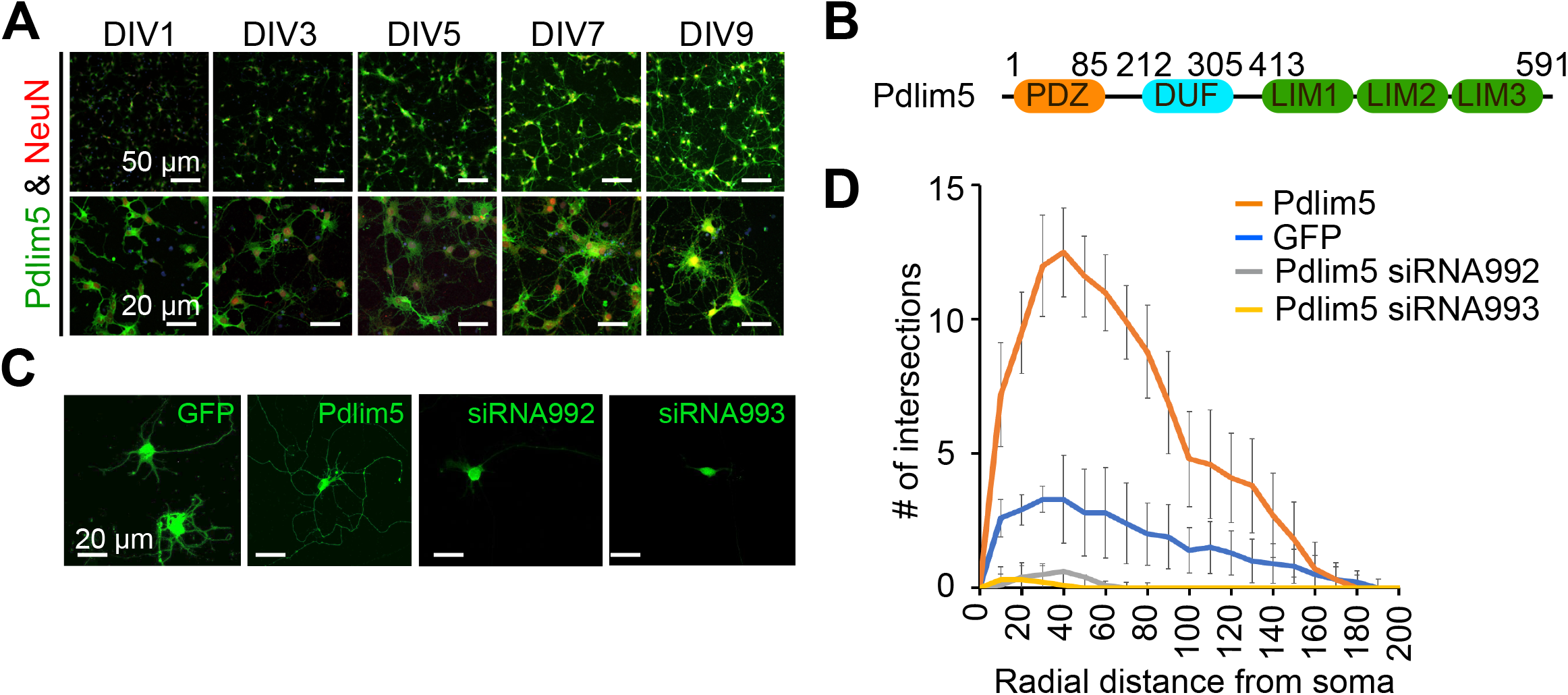
Pdlim5 is a modulator of dendritic branching. A. The images depict rat hippocampal neurons at different developmental stages (DIV1, DIV3, DIV5, DIV7, and DIV9). The neurons are stained for endogenous Pdlim5 (shown in green) and the NeuN marker (shown in red), a marker for neuronal nuclei. Images are representative of 2 biological replicates. B. A diagram illustrating the major subdomains of Pdlim5. The N-terminus contains a PDZ domain (amino acids 1 to 85, shown in orange), followed by a domain of unknown function (DUF, amino acids 212 to 305, shown in cyan), and at the C-terminus, three LIM domains (amino acids 413 to 591, shown in green). C. Immunofluorescent images demonstrating the effects on neuronal morphology of expressing GFP (negative control), Pdlim5, or siRNA constructs targeting the knockdown of Pdlim5. The images reveal a loss of neuronal complexity in cells subjected to Pdlim5 siRNA-mediated knockdown (siRNA992, siRNA993), while exogenous expression of Pdlim5 increases branching. Representative images of three biological replicates. D. Sholl analysis of neurons under the conditions: negative control (GFP), Pdlim5 expression, and Pdlim5 siRNA-mediated knock down. Sholl analysis scores the number of dendrites that intersect a series of evenly spaced concentric rings (generated digitally during data processing) that radiate out from the cell-body/soma. The analysis indicates that compared to GFP-expressing cells (blue line), neurons exogenously expressing Pdlim5 (orange line) exhibited a significant increase in the number of dendrites. Conversely, knockdown of Pdlim5 using siRNA (siRNA992, gray line; siRNA993, yellow line) shows loss of neuronal processes. Data points are the average values (n≥15 neurons) with error bars indicating standard error of the mean (SEM). The significance was assessed using a two-way ANOVA with Bonferroni post-hoc analysis. Scale bars in the images represent 20 µm and 50 µm as indicated.

Here, we reveal that Pdlim5 engages with a key partner protein, PalmD, in dendrite shaping. PalmD is of interest since it appears to be involved in membrane protrusion ^19,20^ necessary to initiate dendrite branching. PalmD is also found to associate with adducin ^20^, a component of the spectrin-actin cytoskeleton that in turn acts in multiple cellular functions ^21,22^. Importantly, as demonstrated in this report, PalmD robustly passed our validation tests for association with Pdlim5, and it has attributes supporting a functional Pdlim5:PalmD relationship in dendrite branching.

## RESULTS

### Pdlim5 is a positive regulator of dendritic branching

In our studies, we have employed mixed cultures of primary rat primary hippocampal neurons in vitro. This selection was made given their faithful reflection of key morphogenic properties found in vivo, while at the same time offering facile manipulation for mechanistic and phenotypic tests. As outlined in Methods, we generally introduce control or experimental constructs into the cultured neurons at DIV1-3 (days in vitro 1-3) and fix at DIV5-7. Here, we assayed for endogenous protein expression as shown for Pdlim5 across DIV1-9. In unperturbed cells Pdlim5 is primarily evident in soma and neurites. It thus appears that in common with PalmD (see below Figure 2), Pdlim5 is available to make potentially-varied contributions across a series of neuronal developmental phases (e.g., from neurite through dendritic stages that are inclusive of initial through advanced branching events and synapse formation) (Figure 1A; see also Figure 2). Further, Pdlim5 and PalmD continue to be expressed in the adult CNS (www.proteinatlas.org). To form a baseline for subsequent phenotypic comparisons, we first built on our earlier findings demonstrating that the exogenous expression of Pdlim5 promotes the branching of rat primary hippocampal dendrites (Figure 1C&D) ^7^. We complemented the exogenous expression phenotypes by conducting Pdlim5 knock-downs. Each of two independent siRNAs efficiently depleted Pdlim5 (Supplemental Figure 1C&D). Further, each siRNA severely reduced branching as seen in representative immunofluorescent images and as quantified by Sholl analysis (Figure 1C&D). Our findings here are entirely consistent with our prior work ^7^ showing that Pdlim5 has a functional role in dendritic branching.

**Figure 2.**
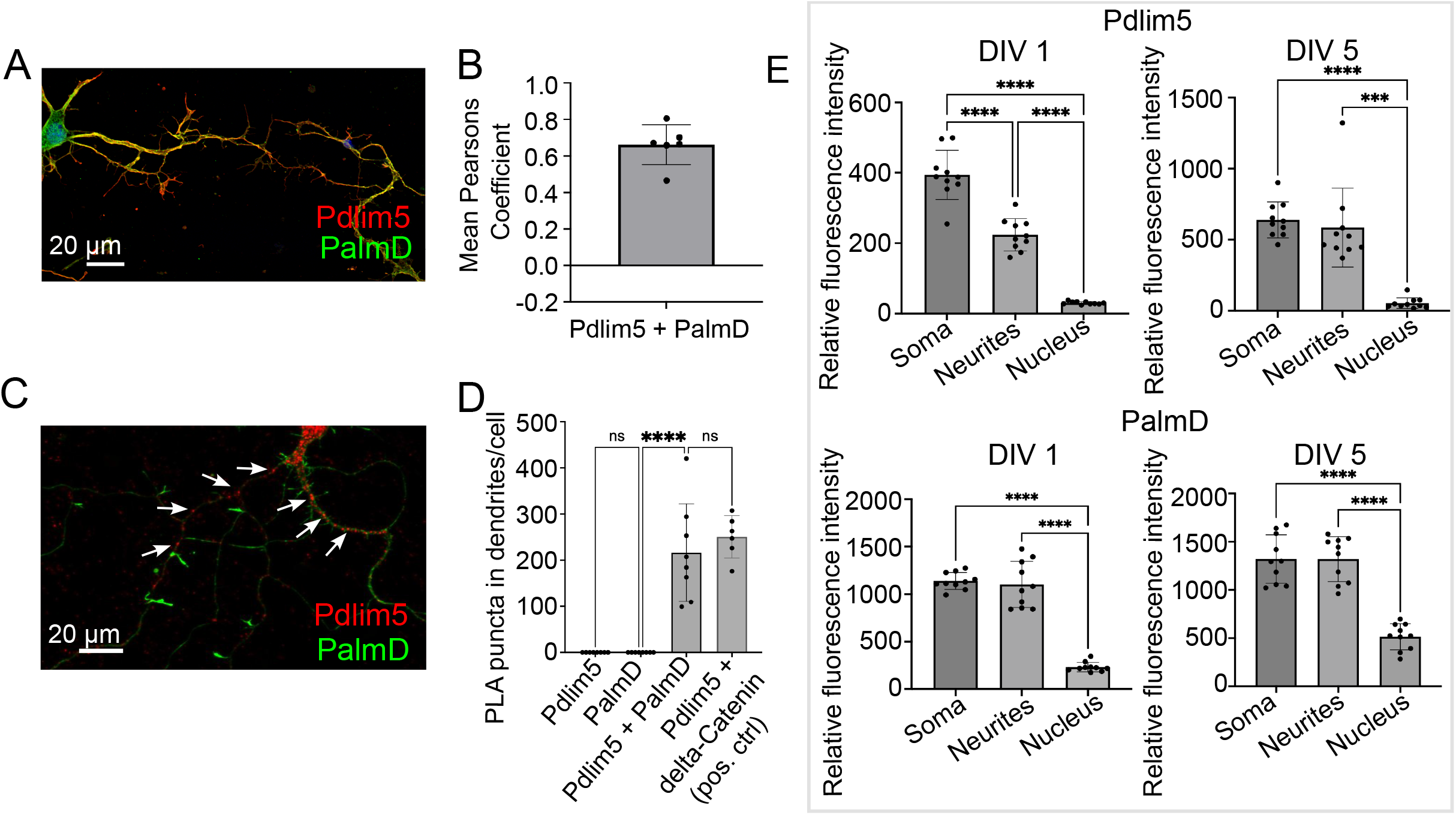
PalmD a novel partner of Pdlim5. A. The image represents the colocalizations of Pdlim5 (shown in red) and PalmD (shown in green) in rat hippocampal neurons at DIV4. Gross colocalization is indicated by the overlap color yellow in regions where both proteins are present within the same cellular compartment. Quantitation in Panel 2B. B. The mean Pearson’s coefficient was calculated to quantify the extent of colocalization between Pdlim5 and PalmD in neurites. The value 0.65 suggests significant (but not complete) linear correlation of Pdlim5 and PalmD colocalization within neurites. Two biological replicates were evaluated with similar outcomes (one shown), with data from ≥5 neurons each. Error bars indicate the standard error of the mean (SEM). Each dot in the graph represents one neuron. C. Immunofluorescent image displays the results of a proximity ligation assay (PLA) of rat hippocampal neurons at DIV4, where the red-colored puncta indicate the positive reaction produced when antibodies to Pdlim5 and PalmD were employed. White arrows point to examples of positive (red) puncta. Quantitation in Panel 2D. D. The quantification of PLA puncta per cell observed in processes is plotted. Antibodies employed against endogenous Pdlim5 or PalmD alone served as negative controls, while the established endogenous complex of Pdlim5:delta-Catenin served as positive control. The data was collected from ≥5 neurons, with each dot representing one neuron. Three biological replicates were evaluated with similar outcomes (one shown). The error bars represent the standard error of the mean (SEM), and the statistical significance, determined using one-way ANOVA, is indicated as P ≤ 0.0001 (****). E. The bar graphs compare the distribution of the Pdlim5 and PalmD proteins in different subcellular compartments, namely soma, neurites, and nucleus, in rat primary hippocampal neurons at two different developmental stages (DIV1 and DIV5). The relative fluorescence intensity for Pdlim5 and PalmD was quantified using ImageJ software. The data was obtained from 10 neurons, with each dot in the bar graph representing data from a single neuron. The error bars indicate the standard error of the mean (SEM). To determine the statistical significance, a one-way ANOVA analysis was performed using GraphPad Prism software. The levels of significance are indicated as **** for P ≤ 0.0001 and *** for P < 0.05, n≥10 neurons. Relative to the nucleus, this signifies high statistical significance of the observed subcellular localizations of both Pdlim5 and PalmD to neurites and soma in rat hippocampal neurons.

### PalmD associates with Pdlim5, likely directly

To glean insight into candidate partners of Pdlim5, we undertook yeast two-hybrid (Y2H) screens of a mouse brain library (Hybrigenics, Inc.). Y2H screens of a chosen “bait” (e.g., Pdlim5) often provide initial clues pointing to direct interactions with “prey” proteins, and ours pointed to PalmD in the company of additional candidates (Supplemental Figure 2). Our interest focused on PalmD from the fact that little is known of its mechanisms of action, yet it has proposed roles in membrane protrusion, an early step in dendrite branching. Further, PalmD is thought to bind (via adducin) to the spectrin-actin cytoskeleton, a significant player in the shaping and functions of neurons ^20-24^. While not pursued here, we note that also arising from our Y2H findings was alpha-actinin4, an established crosslinker of actin microfilaments ^25,26^ that was independently reported to associate with Pdlim5 ^15,16^.

First, we looked at the relative degree of immunofluorescence colocalization of endogenous Pdlim5 and PalmD in rat primary hippocampal neurons (Figure 2A&B). Given that each neuron will go on to bear only one axon, we can assume that we primarily scored neurites destined to become dendritic processes. In DIV4 neurons, colocalization of Pdlim5 and PalmD was evaluated via immunocytochemistry, quantified by employing Mean Pearson’s Coefficient. Values greater than zero are associated with positive correlations. In line with our expectation that Pdlim5 and PalmD form both paired and distinct complexes within cells, our findings are consistent with a significant but not complete overlap of their endogenous localizations.

We next quantitated the relative distributions of Pdlim5 and PalmD in the soma (cellular region immediately surrounding the nucleus), dendritic processes, and nuclei of DIV1&5 rat primary hippocampal neurons (Figure 2E). Given significant areas of co-presence, our findings were consistent with the possibility that these two proteins might form complexes within intracellular neuronal regions undergoing morphological differentiation. However, there remained the need to turn to additional assays for more definitive assessments of direct binding.

In this regard, we applied an approach that is familiar to us, proximity ligation assay (PLA) ^7^. PLA points to very close protein:protein interactions (≤30-40nm) ^7^. In common with our inital Y2H screen, PLA supported the liklihood of a direct Pdlim5:PalmD interaction relative to our negative and positive controls (Figure 2C&D). For example, the Pdlim5:PalmD signals (puncta) observed were comparable in number and intensity to those resolved for our positive-control complex of Pdlim5:delta-catenin ^7^. Overall, our Y2H and PLA findings were supportive of the proposed Pdlim5:PalmD association.

### PalmD associates with Pdlim5 based upon endogenous co-IPs, as well as ectopic Golgi co-relocalization assays

We sought additional validations of the Pdlim5:PalmD association. For endogenous co-immunoprecipitations, we employed rat primary cortical neurons given their relative abundance in comparison to hippocampal neurons. Indeed, the immuno-precipitation of Pdlim5 produced a significant signal for associated PalmD (Figure 3A&B). The inverse co-IP was similarly supportive (Supplemental Figure 3A). Because endogenous Pdlim5 and PalmD migrate via SDS-PAGE at comparable positions, separate blots were used to resolve the co-IPs versus the self-IPs (Supplemental Figure 3A). We additionally assayed co-IPs of exogenous epitope-tagged constructs expressed in HEK293 cells, and we again readily resolved the Pdlim5:PalmD association (Supplemental Figure 3B). Our evidence suggests that the Pdlim5:PalmD association may be quite stable, as we found the complex routinely survived even overnight incubations in cell lysis/extraction buffer.

**Figure 3.**
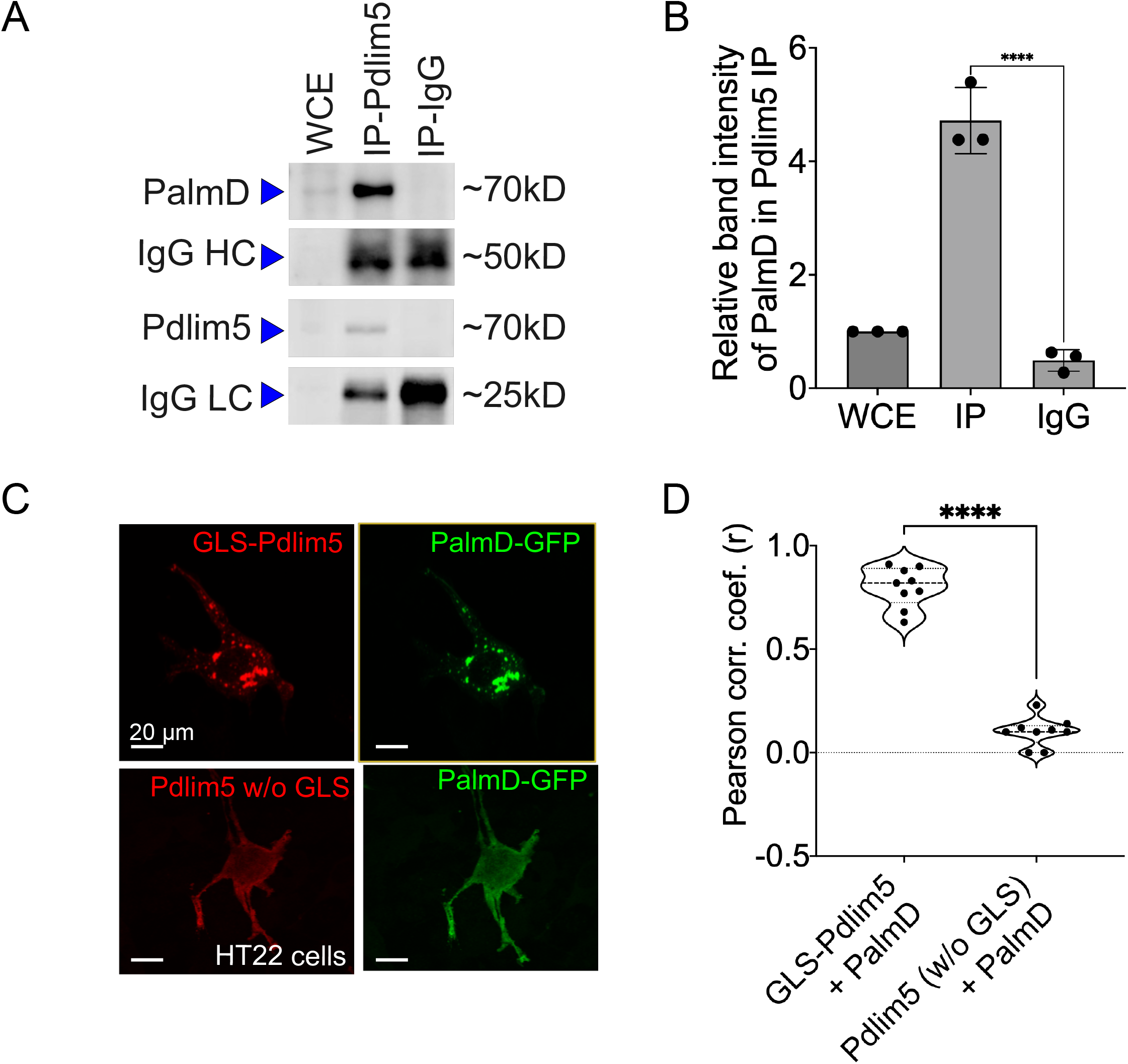
PalmD associates with Pdlim5. A. Immunoblots displaying the endogenous pull-down of Pdlim5 followed by PalmD blotting. Since PalmD and Pdlim5 have similar sizes (approximately 70kD), separate gels were used for blotting of Pdlim5 versus PalmD. n= 3 biological replicates. B. Quantitation by densitometric analysis (Image J software) of co-immunoprecipitations (co-Ips). n= 3 biological replicates of DIV9 rat primary cortical neurons. Each replicate utilized six 10cm dishes containing 2 × 10^6^ cells each (see Methods). The error bars represent the standard error of the mean (SEM), and the statistical significance, determined using one-way ANOVA, is indicated as P ≤ 0.0001 (****). C. Golgi co-relocalization assays demonstrates that Pdlim5 when fused to a Golgi localization sequence (GLS) exhibits an ectopic (red) distribution to the Golgi apparatus as expected. Importantly, PalmD (shown in green) colocalizes with GLS-Pdlim5 to the Golgi. In contrast, when Pdlim5 lacks the GLS, PalmD no longer co-relocalizes with Pdlim5 to the Golgi. n=4 biological replicates, each with analysis of ≥8 cells, scale bar in the images represent 20 μm. HT-22 cells. D. Calculation shows the Pearson’s correlation coefficient for the co-distribution of Pdlim5 and PalmD in the presence versus absence of the GLS-tag upon Pdlim5. The level of statistical significance is indicated as ****, which signifies a p-value of ≤ 0.0001. The violin plot quartiles represent data ranges and middle dark line represent median, and the statistical significance, determined using one-way ANOVA, is indicated as P ≤ 0.0001 (****).

To apply a further test, we took an orthogonal approach that we earlier developed ^7^. In it, one’s construct of interest is ectopically directed to the outer Golgi membrane *via* fusion to a Golgi localization sequence (GLS). A putative partner is then co-expressed, and we score for its ectopic co-targeting, versus not, to the Golgi (we assess if the partner “comes along for the ride”). An attractive aspect of this method is that potential associations are happening in the cell cytoplasm without biochemical treatment for lysis. Thus, the interactions are maintained in a more natural intra-cellular environment. We resolved strong Pdlim5:PalmD colocalization to Golgi when Pdlim5 was ectopically localized there via fusion to a GLS, but not when Pdlim5 lacked such a GLS (negative control) (Figure 3D). These experiments were conducted in HT-22 cells (immortalized mouse hippocampal neuronal cell line), as we found them amenable to Golgi visualization/ scoring. In summary, both our co-IP and GLS tests confirm Pdlim5:PalmD association (Figure 3), as originally supported via yeast 2-hybrid (Y2H) and proximity ligation assays (PLA) (Figure 2).

We went on to apply the GLS methodology to map the region of Pdlim5 that binds to PalmD. Varied constructs of Pdlim5 were tested that retained or lacked each of Pdlim5’s three principal domains: PDZ, DUF, and LIM (Supplemental Figure 3). Our findings clearly indicated a primary role of Pdlim5’s C-terminal LIM domain in PalmD association, as opposed to the N-terminal PDZ or central DUF domains (Supplemental Figure 3D). This result was further supported using traditional co-IP strategies, where the LIM domain was found to co-IP with PalmD (Supplemental Figure 3C).

### Functional assays support Pdlim5’s relationship with PalmD

Analogous to what was earlier indicated in Figures 1A with regards to the developmental presence of Pdlim5 in rat hippocampal neurons (DIV1-9), PalmD is readily observed in dendrites and soma (Figure 4A; see also Figure 2). Using both exogenous expression and knock-down approaches we addressed the impact of manipulating the levels of PalmD in rat primary hippocampal neurons. Upon its exogenous expression, we found that PalmD potently enhanced dendritic branching as quantified via Sholl analysis, while conversely, PalmD knockdown using two independent siRNAs had the opposing impact, dramatically reducing such branching (Figure 4B&C). Similar exogenous expression and knockdown effects were observed when applying other measurements such as counting the number of dendritic tips or branch points per primary hippocampal neuron, or when using Imaris-generated 3-D renderings or binary images for visualizations. (Supplemental Figure 4). This suggested to us that the Pdlim5 and PalmD components within the Pdlim5:PalmD complex are positive modulators of branching.

**Figure 4.**
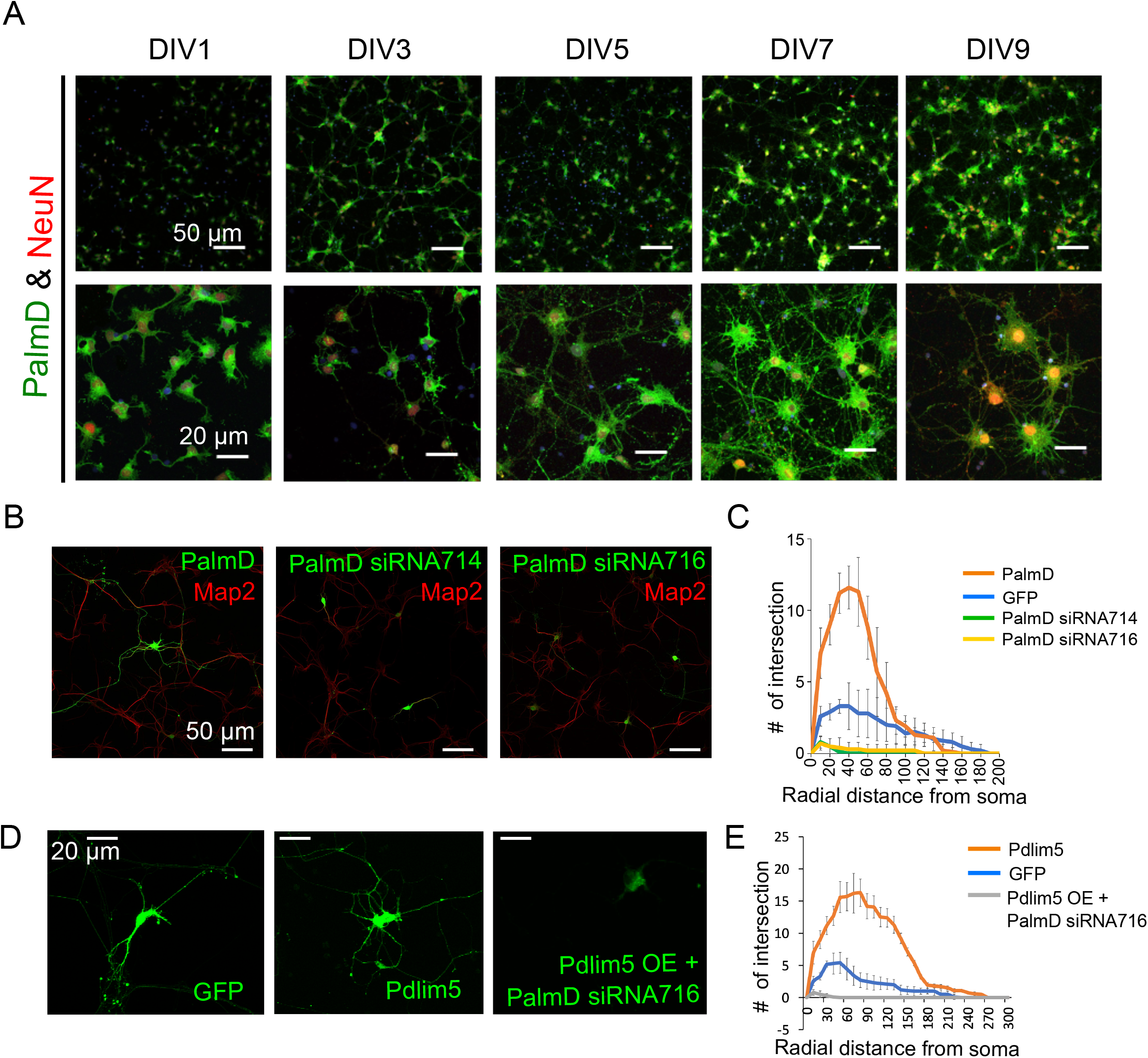
PalmD characterization (?) and its functional dependency with Pdlim5: A. Rat primary hippocampal neurons at the indicated developmental stages (DIV1, DIV3, DIV5, DIV7, and DIV9), are immuno-stained for endogenous PalmD (shown in green) and the NeuN marker (shown in red), which identifies neuronal nuclei. n=2 biological replicates. The scale bars in the images indicate 20 µm and 50 µm. B. These immunofluorescent images illustrate the impact of PalmD exogenous expression versus siRNA-mediated knockdown of PalmD (siRNA714, siRNA716) on neuronal morphology. PalmD siRNA-mediated knockdown decreases process complexity, while exogenous expression increases branching morphology. Three biological replicates in each condition analyzing ≥ 10 neurons. Quantitation present in Panel C. C. Sholl analysis was conducted to examine the dendrite morphology of neurons under different conditions: namely negative control (GFP, blue line), PalmD exogenous expression (orange line), and PalmD siRNA-mediated knock down (siRNA714, green line; siRNA716, yellow line). The analysis demonstrates that neurons overexpressing PalmD exhibit a significant increase in dendritic complexity, while knockdown of PalmD results in severe reductions. The significance was assessed using a two-way ANOVA with Bonferroni post-hoc analysis that varies from P ≤ 0.0001 and P < 0.05, based on radial distance from the cell soma (compared to cells expressing GFP alone). The error bars represent the standard error of the mean (SEM). n≥15 neurons in each of three biological replicates. D. Functional interdependence of Pdlim5 and PalmD. The panel indicates the GFP control condition relative to Pdlim5 exogenous expression, and versus PalmD exogenous expression in the presence of PalmD siRNA-mediated knockdown (KD). The results suggest an interdependence between Pdlim5 and PalmD in regulating neuronal morphology. Analyzed ≥ 10 neurons in each of three biological replicates. See panel E for quantitation. E. Sholl analysis was performed to analyze dendrite morphology under different conditions: GFP control, Pdlim5 exogenous expression, and Pdlim5 exogenous expression with concomitant knock down of PalmD (siRNA716). The analysis reveals that neurons overexpressing Pdlim5 (orange line) exhibit increased numbers of dendrites compared to control GFP-expressing cells (blue line) (P ≤ 0.0001 and P < 0.05 for the respective regions 20-90 µm and 110-170 µm from the soma). However, when PalmD siRNA716 is introduced alongside Pdlim5 exogenous expression (gray line), there is a significant loss of branching function. The error bars represent the standard error of the mean (SEM). n≥15 neurons each from three biological replicates. The significance was assessed using a two-way ANOVA with Bonferroni post-hoc analysis.

We next asked if the effects observed upon expressing Pdlim5 was dependent on the presence of PalmD, and vice versa. When Pdlim5 was exogenously expressed under conditions of PalmD knockdown, we failed to observe any enhancement of branching (Figure 4D&E). Indeed, as seen with the knockdown of PalmD-alone, we observed suppression of branching relative to the GFP-alone control. We likewise observed that the promotion of dendrite branching upon PalmD expression became suppressed upon the parallel knockdown of Pdlim5 (Supplemental Figure 4E&F). These findings are consistent with the existence of not only a biochemical Pdlim5:PalmD complex but also a functional dependence between Pdlim5 and PalmD in the process of dendritic branching.

## DISCUSSION

The complexity of neuronal interconnections allowing for vertebrate central nervous system functions is impressive by any standard, and there remains much to be learned about how dendrites are shaped to enable such interconnections. Generally, neuronal networking requires the control of dendrite morphology because neuron:neuron interactions depend upon the precise postsynaptic contacts of each dendrite’s associated synaptic spines. The postsynaptic specializations of dendrites receive signals from the presynaptic terminals of axons that extend from nearby or distal neurons. In this report, we focused upon the gross shaping of dendrites as opposed to the formation of precise synaptic contacts, but these vastly different morphologic scales remain interrelated given that proper synapse patterning can only come about when dendrites adopt morphologies consistent with spine/ synapse generation. There are numerous extrinsic and intrinsic cellular factors that modulate neuronal branching. With regards to extrinsic cues, glutamate ^27,28^ and brain derived neurotrophic factor (BDNF) ^29^ are just two examples, along with varied transmembrane ligand:receptor pairings inclusive of cell:cell adhesion molecules ^30,31^. Prominent intrinsic cellular factors include examples like small-GTPases, cytoskeletal components or regulators (e.g., affecting organelle distribution and the extent or polarity of vesicular traffic), and varied intracellular or junctional signaling entities ^32^. Nuclear contributions (e.g., transcriptional) are also relevant in establishing gross or refined morphologies but their varied effects are generally viewed as being more indirect.

Using rat primary hippocampal neurons, we here reveal that increasing PalmD levels heightens dendritic branching, while decreasing PalmD has the opposite effect. PalmD has been suggested to engage in shaping membranes, potentially by enhancing their curvature while facilitating intracellular interactions with actin-based processes ^19,20^. Such a possibility is consistent with properties ascribed to related family members of PalmD such as paralemmin ^33-36^, for which C-terminal palmitoylation and thereby enhanced membrane localization was indicated to promote neuronal process complexity. Recently, PalmD was reported to promote process formation in the basal progenitor cells of the mammalian neocortex ^20^. In that work, PalmD bearing a C-terminal Caax motif was proposed to achieve membrane localization via Caax palmitoylation in line with its activity in process formation. Our findings were instead derived from differentiated rat hippocampal and cortical neurons. They indicate that even a PalmD isoform lacking the C-terminal Caax motif (isoform having four terminal KKVI residues instead ^20,37^) remained robustly capable of advancing dendritic branching. In the neuronal context, one possibility is that PalmD lacking its C-terminal Caax motif might still become palmitoylated or otherwise lipidated elsewhere within its structure. This could enable PalmD’s presence and interactions at presumptive or forming dendritic membranes to advance branching processes. Alternatively, a fraction of PalmD might become membrane localized through its direct association with Pdlim5 or other potential membrane associated binding partners. Membrane localization was artifically reflected here using an experimental approach we earlier devised (GLS assay) ^7^. In it, we targeted Pdlim5 to the Golgi and demonstrated co-targeting of PalmD, that is, Palm D came along for the ride. In normal neuronal contexts, the PDZ-domain of Pdlim5 is capable of binding to multiple proteins bearing PDZ-motifs, presumably including that proportion of delta-catenin at dendritic membranes ^7^. Other possibilities to consider as tethers to the membrane-cytoskeleton include adducin ^20^, as well as potentially indirect interactions of PalmD with alpha-actinin or cortactin (see below). In all cases, while beyond the scope of our report here, future work in the field is required to address PalmD’s downstream mechanisms of action, inclusive of its indirect associations with the inner dendritic membrane or with relevant cytoskeletal elements.

Using multiple orthogonal tests, we provide evidence of PalmD’s association with Pdlim5, a scaffold that we earlier characterized in association with delta-catenin ^7^. Each of these three proteins enhance dendrite branching when exogenously expressed alone, and conversely, the knockdown of each individual gene product alone leads to neurons having a greatly reduced presence of dendritic processes. delta-Catenin is thought to promote dendrite branching in part through inhibition of the small-GTPase RhoA ^38-41^, which we have likewise indicated is the case for Pdlim5 ^7^. Given that PalmD associates with Pdlim5, there is the possibility that PalmD’s effects may likewise involve RhoA or another small-GTPase. Further potential links of PalmD to cytoskeletal modulation revolve around the presence of additional associated proteins in complex with Pdlim5 or with delta-catenin. For example, associations based upon the respective interactions of Plim5 and delta-catenin with the potent cytoskeletal regulators alpha-actinin ^15,16^ and cortactin ^40,41^, each known to contribute to the production or maintenance of cellular processes. Thus, while conjectural at this stage, a number of possible downstream mechanisms of action could be proposed based upon established associations of PalmD’s direct or indirect protein partners.

Our prior work ^7^ and that of others ^41^ outlines what could be a key upstream pathway that impinges on the Pdlim5:PalmD complex revealed here, with consequent effects upon gross dendritic morphology. Specifically, Pdlim5’s (and thereby PalmD’s) association with delta-catenin is determined by a “phospho-switch” at the very C-terminus of delta-catenin. As reflected in this report’s Graphical Abstract, the phosphorylation of delta-catenin is responsive to upstream glutamate signaling via the neuronal mGluR5 receptor and consequent activation of Cdk5 kinase ^7^. Even as future work will be needed to precisely identify the cytoskeletal players downstream of the delta-catenin:Pdlim5:PalmD complex, we speculate as noted the direct involvement of cortactin, alpha-actinin4, and RhoA inhibition, as well as the spectrin network and membrane-curvature agents. In summary, we unveil here a novel Pdlim5:PalmD complex that associates with delta-catenin, which through functional interdependencies, promotes dendritic branching-morphology required in development and is aberrant (see Introduction) in multiple pathological contexts.

## Supporting information

Supplemental Figure 1. Pdlim5 conservation and function

Supplemental Figure 2. Selected Pdlim5 associated candidates with known cytoskeletal functions

Supplemental Figure 4. Function and functional dependency of Pdlim5: PalmD complex

## ACKNOWLEDGEMENTS

We thank past lab member Ryan Baumert PhD for being instrumental in initiating our group’s work upon dendrite morphology ^7^. For helpful discussion we thank the following primary investigators and the members of their labs: Malgosia Kloc PhD, Rachel Miller PhD, and Jaeil Park PhD. Use of the A1-Nikon (confocal images) was made possible via the UT MDACC Department of Genetics NIH Instrumentation Grant 1S10OD024976-01 (AP). Assistance with DNA sequencing was provided from the National Cancer Institute Core Grant CA-16672 to UT MDACC. This work was supported by NIH 1RO1MH115717 (PDM & MNW). MNW acknowledges an endowment from the William Wheless III Professorship, and PDM acknowledges an Ashbel Smith Professorship.

The authors declare no competing financial interests.

## AUTHOR CONTRIBUTIONS

YS, PDM and MNW planned the project. YS analyzed the data in communication with PM and MNW. YS conducted the bulk of the experiments, with LLM and MNW preparing the rat primary hippocampal and cortical cultures. MD undertook the proximity ligation assays of Pdlim5 with PalmD, and ascertained their endogenous localizations via immunofluorescence, with assistance from LLM and MNW. APH assisted with advising YS and the laboratory on imaging. The manuscript was written by YS and PDM with consistent input from MNW. All authors read and approved submission of the final version of the manuscript.

## METHODS

### Neuronal cultures and transfection

Primary hippocampal and cortical neurons were obtained from rat embryos at the 18th day of gestation (E18), following methodology previously described in ^42,43^. Subsequent to isolation, the hippocampal neurons were plated in 24-well tissue-culture plates at a density of 2 × 10^5^ cells per well. To facilitate adhesion, glass coverslips within the wells were pre-coated with 100 µg/ml poly-D-lysine (Sigma-Aldrich). Cortical neurons were plated at 20 × 10^6^ cells per 10 cm Petri dish pre-coated with 100 ug/ml poly-D-lysine. The cultured hippocampal neurons were maintained in Neurobasal Medium (Life Technologies), supplemented with B-27, GlutaMAX, and penicillin-streptomycin (each from Life Technologies). At 3 days in vitro (DIV), the neurons underwent Lipofectamine 2000 based transfection using the manufacturer’s protocol (Life Technologies) but employing 1 µg plasmid DNA.

### HEK293 cell culture and transfection

HEK293 cells from ATCC (ATCC# CRL-1573) cells were cultured in DMEM cell culture media (Sigma-Aldrich) supplemented with 10% FBS (Sigma-Aldrich) and 1x penicillin-streptomycin (Life Technologies). Media change was performed every other day until the cells reached 80% confluency, after which they were transferred to new culture dishes. Transfection of the HEK293 cells with exogenous DNA (1 µg) was accomplished using Lipofectamine 2000 as per the manufacturer’s instructions. Transfection took place when the cells reached 50-60% confluency in six-well culture plates. Co-IPs were conducted as below mentioned.

### cDNA constructs

All cDNA constructs were cloned into the backbone of the pCS2 mammalian expression vector. Engineering was undertaken by Epoch Life Science and/ or the McCrea laboratory. All constructs were confirmed with DNA sequencing. The full length PalmD-pCMV6-AC-GFP was purchased from Origene (RG203079), and PalmD-GFP was moved into the pCS2 vector. Pdlim5 was engineered previously in our laboratory ^7^. DNA maxi-preps were outsourced to Epoch Life Sciences. See STAR Methods Key Resources Table for a complete list of constructs and their corresponding epitope tags/ fusions.

### siRNAs

To synthesize and validate siRNAs that target Pdlim5 or PalmD, we first identified corresponding 19-nucleotide target sequences using the siRNA Wizard v3.1 tool from InvivoGen. Two siRNAs were selected for each target to enable the generation of siRNAs. The siRNA sequences for Pdlim5 were 5ʹ-GCACUGUAUUGUGAGCUAUtt-3ʹ and 5’-CAACUGUGCUCACUGCAAAtt-3’, while for PalmD they were 5’-GCAUCAGGCAGAACGAAUAtt-3’ and 5’-CGAGGAUAUCUAUGCUAAUtt-3’.

siRNAs for Pdlim5 and PalmD were purchased from Life Technologies. Individual siRNAs were transfected into hippocampal neurons using the manufacturer’s Lipofectamine 2000 protocol. Neurons were allowed to grow for 24-48 hours before conducting immunofluorescence assays. Knockdown efficiency of each siRNA was assessed using immunoctyochemistry with anti-Pdlim5 or anti-PalmD primary antibodies, and IMARIS software was used to quantify the intensity of immunostained images. Sholl analysis was used to assess the impact upon neuronal morphology (see below).

### Antibodies

Antibodies that recognize specific epitope tags as well as endogenous proteins were obtained commercially. Rabbit polyclonal anti-Myc epitope-tag (MT) antibodies (Cell Signaling Technology/ CST #2272S), mouse monoclonal anti-Myc epitope-tag (MT) antibodies (CST #9B11), anti-GFP antibodies (CST #2956 from rabbit and CST #4B10 from mouse), and a mouse monoclonal anti-FLAG epitope-tag antibody (Sigma #F7425). For the detection of endogenous Pdlim5, two mouse monoclonal antibodies were utilized: one for the purpose of IP/ immuno-blotting (Thermo Fisher Scientific catalog # MA5-25915) (Figure 3A and S3A), with a second mouse monoclonal (Sigma-Aldrich catalog # WH0010611M1) used to detect endogenous Pdlim5 via immunofluorescence (Figure 1A). A rabbit polyclonal antibody directed against Pdlim5 (#38–8800) was used for both blot and immunofluorescence (IF) assays. Primary rabbit IgG from Life Technologies (#10500C) was used for negative-control “IPs”, as was mouse IgG from Invitrogen (#10400C). (Figure 3A, S3A, S3B, S3C). To detect endogenous PalmD in immunofluorescence assays, we employed a rabbit polyclonal antibody (ProteinTech, catalog # 16531-1-AP) (Figure3A and S3A). delta-Catenin was visualized using a mouse monoclonal antibody (BD Transduction Laboratories catalog # 611537, not shown). All HRP-conjugated goat polyclonal secondary antibodies were obtained from Thermo Fisher Scientific (anti-mouse #31430 and anti-rabbit #31460). All Alexa fluor immunofluorescent secondary antibodies were purchased from Invitrogen (488 anti-mouse #A32723, 488 anti-rabbit #A22008, 555 anti-mouse #A21422, 555 anti-rabbit #A32732 and 555 anti-chicken A32932).

### Golgi Localization Sequence Assay (GLS)

Immortalized mouse hippocampal neuronal cells (HT-22) were cultured in DMEM cell culture media (Sigma-Aldrich) supplemented with 10% FBS (Sigma-Aldrich) and 1x penicillin-streptomycin (Life Technologies). Transfections of cells were done at 50%-to-60% confluency using Lipofectamine 2000. Golgi co-relocalization was examined by fixation with 4% PFA 24 hours after transfection, followed by immunostaining and subsequent image collection using confocal microscopy. To ectopically relocalize Pdlim5 to the Golgi, Pdlim5 was tagged at its’ N-terminus with the Golgi localization sequence (GLS) of mammalian target of rapamycin (mTOR) ^7^. Following co-transfection with one of Pdlim5’s putative partners (e.g., PalmD), the cells were subjected to immunofluorescence staining as previously described ^7,44^. Qualitative analysis involved visual examination of co-relocalized partner proteins to the Golgi, occurring only when GLS-Pdlim5 is present. Quantitative analysis involved measuring the average fluorescence intensity of the Golgi body compared to that of the cytosol in each optical channel using ImageJ. This comparison generated a Golgi:cytosolic intensity ratio for each condition.

GLS from mammalian target of rapamycin: 5ʹ-ACTAGTGAAATGCTGGTCAACATGGGAAACTTGCCTCTGGATGAGTTCTACCCAGCTGTGTCCATGGTGGCCCTGATGCGGATCTTCCGAGACCAGTCACTCTCTCATCATCACACCATGGTTGTCCAGGCCATCACCTTCATCTTCAAGTCCCTGGGACTCAAATGTGTGCAGTTCCTGCCCCAGGTCATGCCCACGTTCCTTAACGTCATTCGAGTCTGTGATGGGGCCATCCGGGAATTTTTGTTCCAGCAGCTGGGAATGTTGGTGTCCTTTGTGAAGAGCCACATCAGACCTTATATGGATGAAATAGTCACCCTCATGAGA-3ʹ.

### Proximity Ligation Assay (PLA)

To perform Proximity Ligation Assays (PLA), neurons were fixed at 4 days in vitro (DIV4) with 4% PFA and the Duolink PLA kit was employed as previously described ^7^ following the manufacturer’s (Sigma-Aldrich) protocol ^45^.

In brief, cells were permeabilized with 0.5% Triton-X 100 and blocked with 1% BSA blocking solution. Neurons were then incubated with primary antibodies specific to each of the two endogenous target proteins, such as Pdlim5 and PalmD. Subsequently, the cells were washed with PBS and incubated with Duolink PLA probes. These probes recognize the primary antibodies and enable the generation of a DNA structure recognized by manufacturer-provided fluorescent probes. To generate the DNA structure, the cells were incubated with a ligase and amplification enzymes. Finally, the permeabilized and blocked cells were incubated with the fluorescent probes and imaged using a Nikon T2i Inverted Confocal Microscope equipped with an Apo-Plan 60× 1.4 NA oil objective. The imaging was conducted at a pixel size of 100 nm, and image stacks were converted into maximum projection images for analysis. The quantification of puncta was performed using the Puncta Analyzer plugin within the ImageJ software.

### Co-immunoprecipitations and immunoblots

For immunoblot assay of endogenous proteins from rat primary cortical neurons, 2 × 10^6^ cells from six 10 cm dishes were collected and lysed at DIV9. For immunoblotting assays of exogenous co-immunoprecipitated proteins, HEK293 cells were lysed 24-to-48 hours following their co-transfection, using established protocols ^7^. In brief, following lysis and pelleting of debris, protein concentrations of lysates were equalized (500-1000 µg per condition/ IP) and incubated with primary antibodies for 2-to-12 hours at 4°C with gentle rotation. Protein-A and Protein-G Dynabeads (Invitrogen, #10002D and #10004D) were added to each tube and incubated for an additional 20-30 min with gentle rotation at 4°C. After washing, the associated proteins were eluted from the beads upon addition of 2× sample buffer with βME and heated at 95°C for 5 min. Co-IP samples, or in the case of whole-cell extracts equalized cell lysates, were then loaded onto 10% polyacrylamide gels using a BioRad mini-PROTEAN Tetra Cells, 2-gel system setup (#1658005) and run for 120 minutes. Proteins were then transferred to nitrocellulose membranes (GE Whatman) using a Pierce Power Blotter (Thermo Fisher Scientific). Membranes were blocked in TBS supplemented with 0.1% Tween-20 plus 1% dry milk, with incubation for 4-to-12 hours at 4°C. Blocked membranes were incubated in primary antibodies overnight at 4°C before being incubated with HRP-conjugated secondary antibodies (Invitrogen) for 1-to-2 hours at room temperature. Lastly, membranes were incubated with Pierce ECL immuno-blotting substrate (Thermo Fisher Scientific # 32106) and imaged using a ChemiDoc MP imaging system (Bio-Rad).

### Immunofluorescence staining and Confocal microscopy

Immunostaining of rat primary hippocampal neurons or HT-22 cells employed established protocols (Baumert at al., 2020; Lee et al., 2014). Briefly, cells cultured on poly-D-lysine-coated coverslips were fixed with 4% paraformaldehyde for 10 minutes at room temperature and then permeabilized with 0.5% Triton X-100 solution for 15 min. Following fixation and permeabilization, cells were blocked with PBS containing 1% bovine serum albumin overnight at 4°C. Cells were then incubated with the indicated primary antibodies overnight at 4°C and subsequently rinsed three times with fresh PBS for 10 min each, followed by an overnight rinse with PBS at 4°C. Cells were then incubated with Alexa Fluor fluorescent dye-conjugated secondary antibodies (Invitrogen) for 1 hour at room temperature before the coverslips were washed and finally mounted onto glass slides with Vectashield (Vector Laboratories) mounting solution. Cells were visualized using a Nikon T2i Inverted Confocal Microscope with an Apo-Plan 60× 1.4 NA oil objective at room temperature. Images were captured with a Nikon A1-DUG GaAsP hybrid four-channel multi-detector and Nikon NIS-Elements software. All z-series images were acquired at a pixel size of 100 nm and a step size of 0.2 µm.

### Neuronal morphological analysis by IMARIS9.9

Rat primary hippocampal neurons underwent assessment of their morphological characteristics with a focus upon neurites/ dendrites. z-Series images of immunofluorescently stained neurons were captured using a Nikon T2i Inverted Confocal Microscope, as previously mentioned. Prior to analysis, IMARIS 9.9 was utilized to generate a 2-dimensional maximum-projection image with background subtraction from the confocal z-series of each neuron. In cases where dendrite number was evaluated, counting focused upon the number of dendrite tips per neuron, excluding any protrusions shorter than 5 µm. Dendritic length was measured by tracing from the cell body to the tip of the dendrite using IMARIS 9.9 filament tracer. Thresholds and settings for IMARIS 9.9 filament tracer were kept the same for all samples. For Sholl analysis, the maximum-projection images were converted to binary and analyzed in IMARIS 9.9 filament tracer using concentric rings with a 5 µm step size to evaluate dendrite morphology in relation to the distance from the cell body ^44^.

### Statistical analysis

Data were analyzed using GraphPad Prism. Statistical significance for Sholl data were determined using a two-way ANOVA with Bonferroni post-hoc analysis (figure 1D, 4C, 4E and S4E). For all other comparisons, a one-way ANOVA with Tukey’s test was used (Figure S1B, S1D, 2B, 2D, 2E, 3B, 3D, S4B, S4C and S4D). Data distribution was assumed to be normal, but this was not formally tested. Significance was assigned at P < 0.05. Please refer to each Legend for specific information related to experimental statistical considerations.

## SUPPLEMENTAL FIGURE TITLES & LEGENDS

**Supplemental Figure 1.**
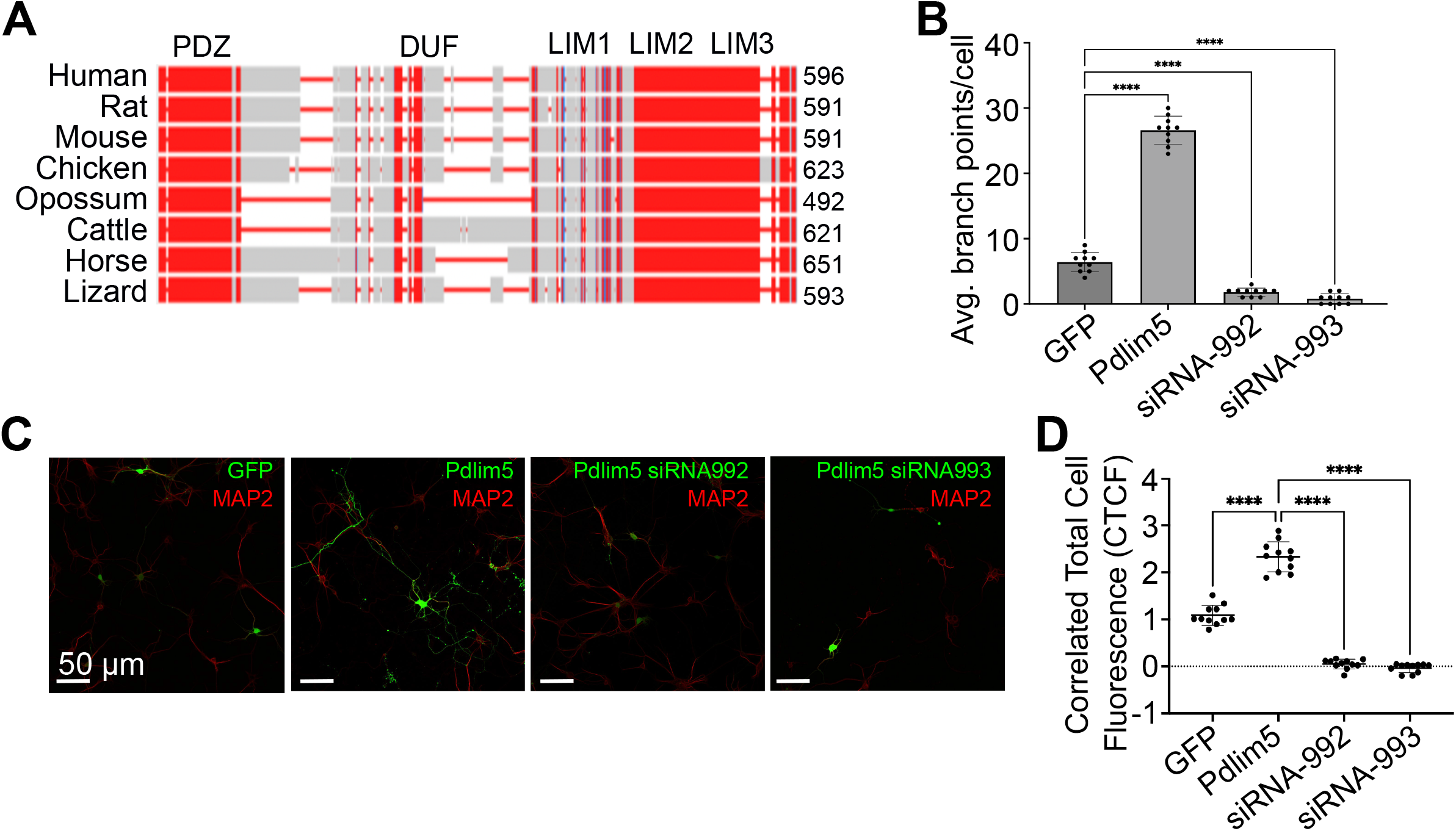
Pdlim5 conservation and function. A. The conservation of Pdlim5 among various vertebrates is depicted in the figure. Conserved sequences are represented in red, while identical sequences are shown in gray. Gaps in the alignment are denoted by red lines. The number of amino acids for each species is indicated in the top right corner, and the species names are listed on the left side. The multiple sequence alignment was conducted using Clustal Omega software. B. Impact of Pdlim5 knockdown (siRNA992, siRNA993) on hippocampal neuron branching morphology. Exogenous expression of GFP (control), Pdlim5, and siRNA-mediated knockdown of Pdlim5. Following siRNA-mediated knockdown of Pdlim5 (siRNA992, siRNA993), we observed a significant reduction in dendritic complexity. In contrast, the exogenous expression of Pdlim5 resulted in an increase in branching morphology in comparison to negative control GFP. n=10 neurons each from three biological replicates. C. Illustrative images demonstrating the knockdown (KD) of Pdlim5. The control GFP image (green, first image on the left) exhibits less branching compared to the expression of exogenous Pdlim5 (green, second image from the left), while loss of branching morphology is observed in the Pdlim5 knockdown siRNA images (green, third and fourth images from the left). Neuron health was assessed using the Map2 marker. n = 3 biological replicates. The scale bars in the images indicate 20 µm. See panel D for quantitation. D. The Pdlim5 total cell fluorescence was quantified using ImageJ for negative control (GFP), positive control Pdlim5, and siRNA mediated knockdown of Pdlim5 (siRNA992, siRNA993). As expected in comparison to control, exogenous Pdlim5 significantly increases Pdlim5 total cell fluorescence while knockdown significantly decreases Pdlim5 total cell fluorescence. n≥10 neurons in each of three biological replicates. The quartiles represent data ranges and middle dark line represent median, and the statistical significance, determined using one-way ANOVA, is indicated as P ≤ 0.0001 (****).

**Supplemental Figure 2.**
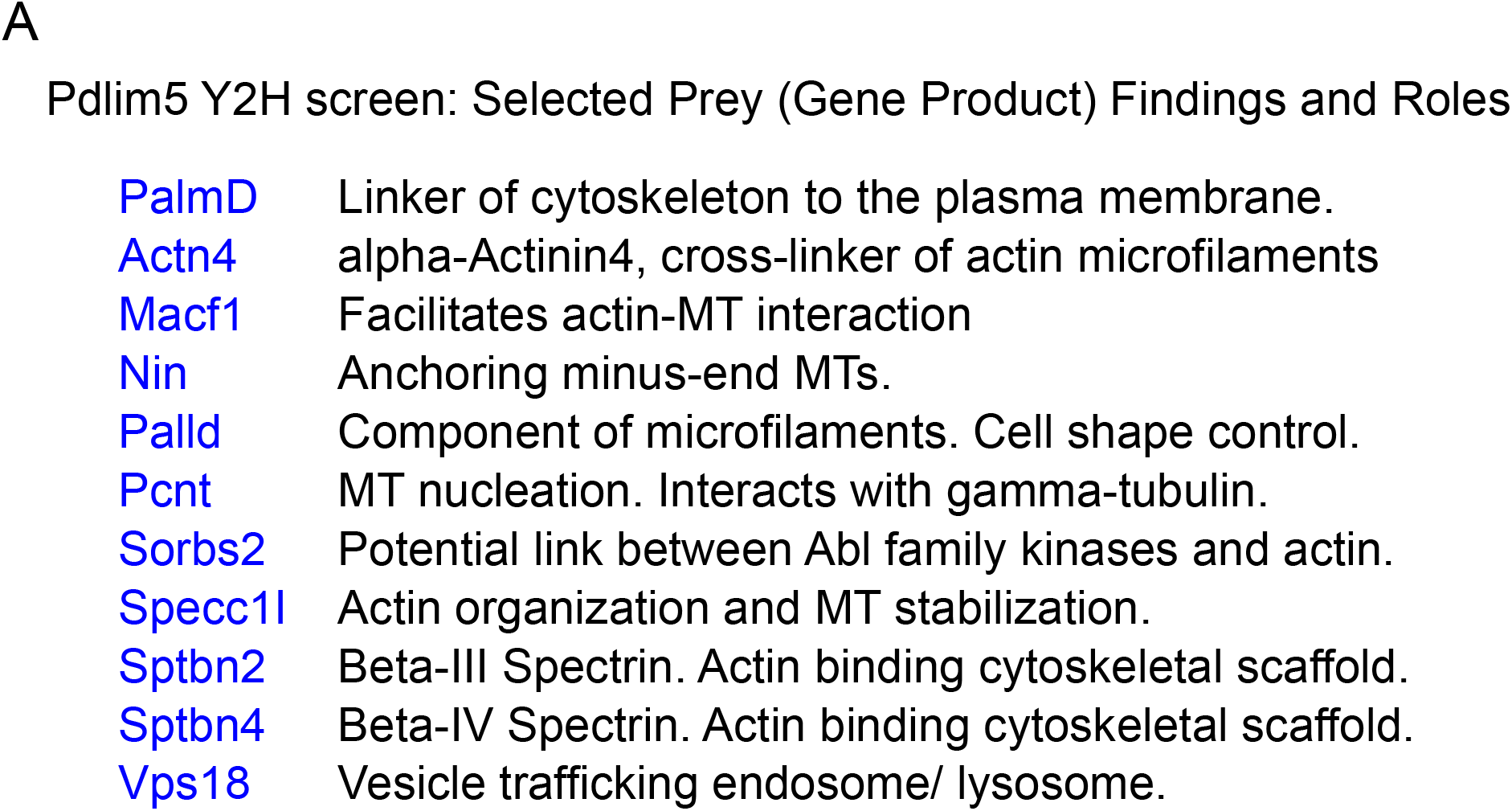
Selected Pdlim5 associated candidates with known cytoskeletal functions. Our yeast-2-hybrid (Y2H) screen suggested several Pdlim5-associated candidates involved in cytoskeletal functions. In total we resolved over 70 candidates with 11 being implicated in cytoskeletal contexts, including for example Beta-III Spectrin, Beta-IV Spectrin, and alpha-Actinin4. Our studies here focused upon PalmD for reasons outlined in the text.

**Supplemental Figure 3.**
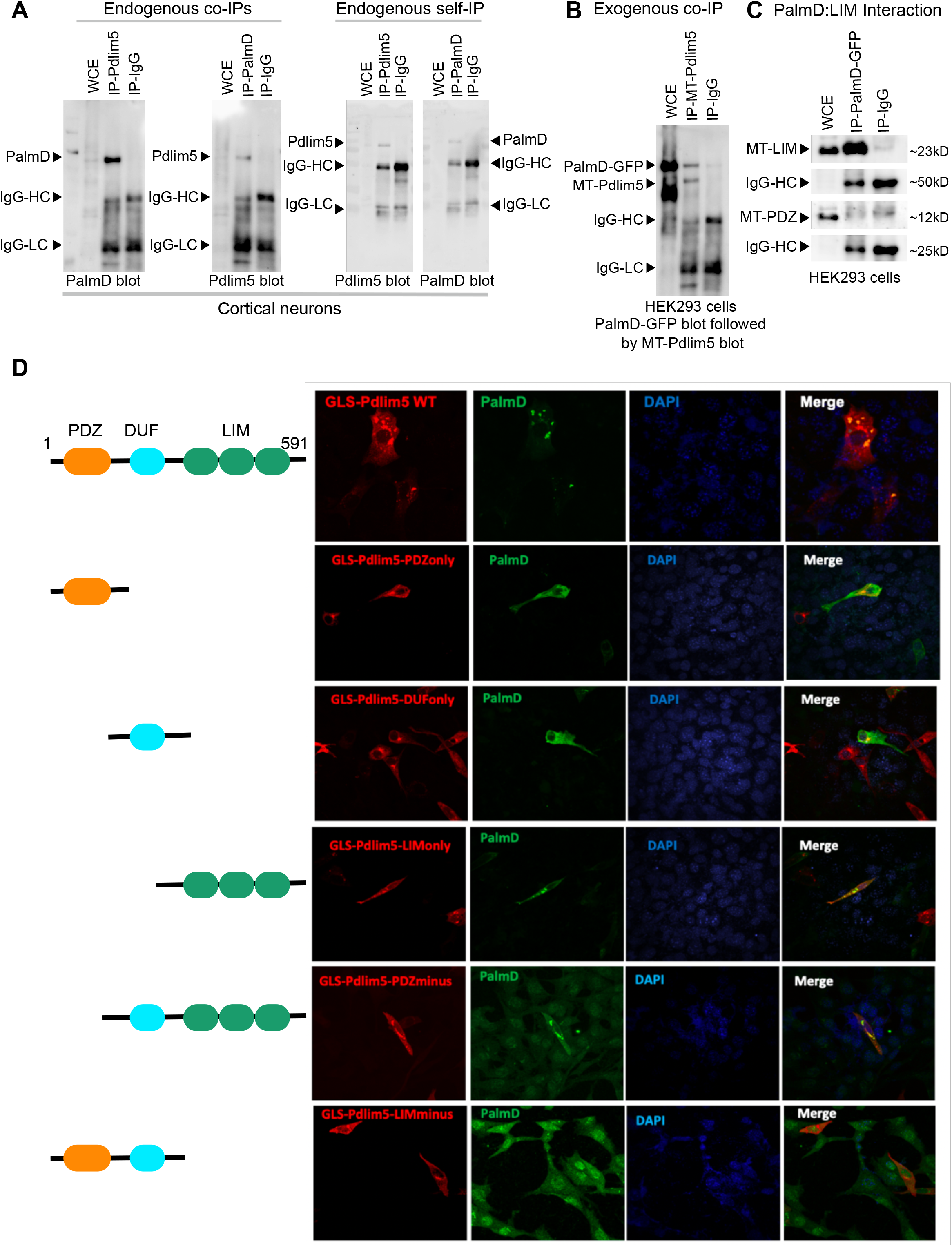
PalmD, a novel partner of Pdlim5, associates with Pdlim5’s LIM domain. A. The endogenous co-immunoprecipitation (co-IP) of Pdlim5 and PalmD was performed from rat primary cortical neurons at DIV9. Blots were used to demonstrate the pull-downs of Pdlim5 and PalmD. Separate blots for endogenous PalmD and Pdlim5 were undertaken given their similar molecular weights (∼70kDa). Endogenous self-IP blots were also performed, confirming the effectiveness of the endogenous IP procedures. n≥3 biological replicates for each condition. B. In exogenous contexts (HEK293 cells), we engineered Pdlim5 with a 6x-Myc-epitope tag (MT) at its N-terminus (∼75kD), while PalmD was fused with GFP (∼100kD). Together with the findings noted in Figures 2 and 3, and immediately above in Panel A, we collectively provide evidence of both endogenous and exogenous association between Pdlim5 and PalmD. n≥3 biological replicates in each condition. C. PalmD co-immunoprecipitation with Pdlim5’s isolated LIM domain. PalmD was co-expressed with each individually isolated domains of Pdlim5 (LIM or PDZ). The results revealed that PalmD associates specifically with the LIM domain of Pdlim5. Relative to negative controls, no association was observed with the other isolated domains, such as PDZ (or with the central DUF domain; not shown). n≥3 biological replicates in each condition. See also below Panel D. D. In addition to using co-IP (see panel C above), we applied an orthogonal method we term GLS (see the text and Methods), to explore the association of different domains of Pdlim5 with PalmD. The assay revealed the colocalizations of PalmD with Pdlim5 at the Golgi only when the LIM domain of Pdlim5 was present. This finding again suggests that PalmD has an association with the LIM domain of Pdlim5. n≥3 biological replicates.

**Supplemental Figure 4.**
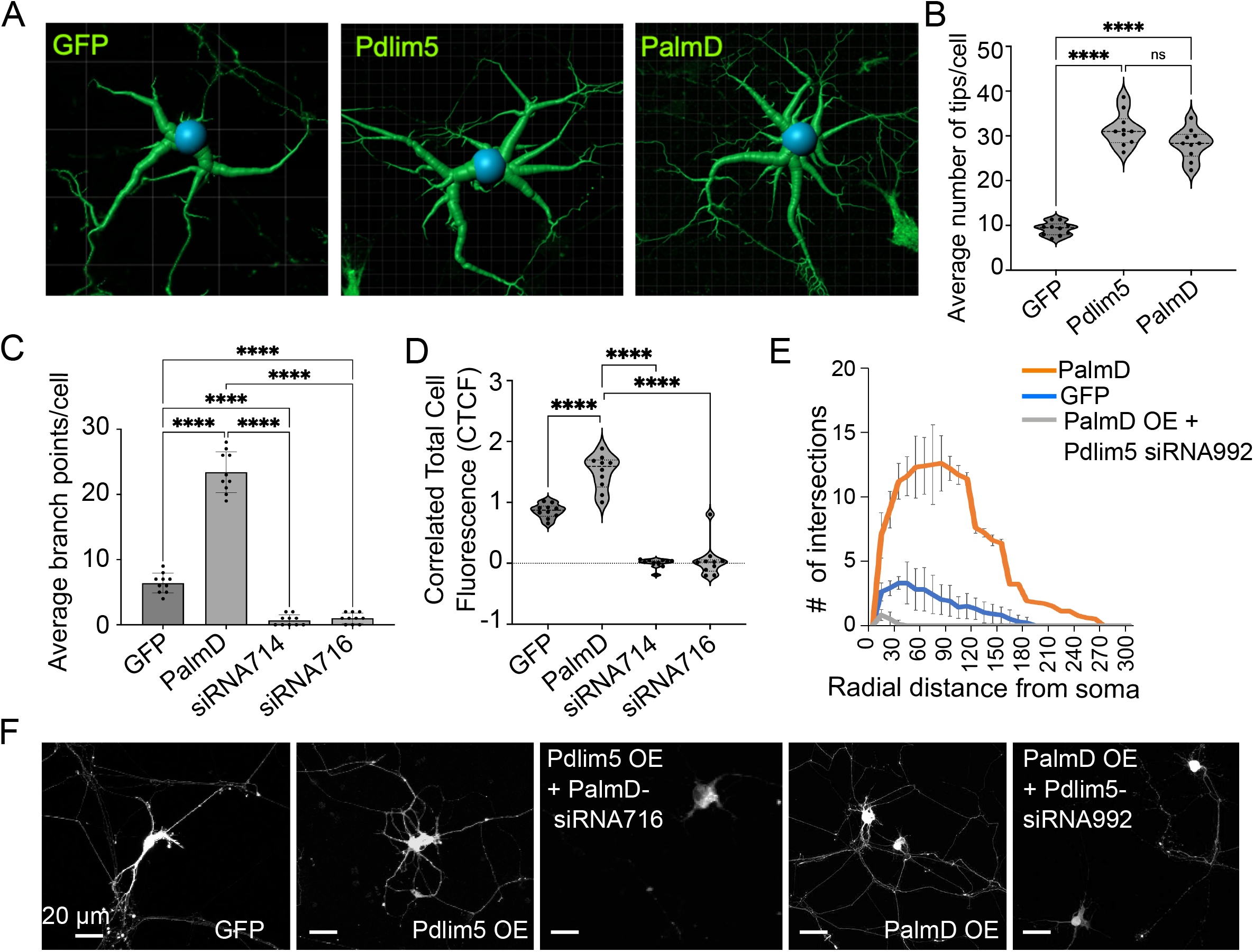
Function and functional dependency of Pdlim5: PalmD complex. A. Imaris-generated 3D rendering image of rat primary hippocampal neurons shows the exogenous expression of negative control GFP, Pdlim5, or PalmD. Increased branching is observed in neurons expressing Pdlim5 or PalmD compared to the GFP control. Panel B provides quantitation of the average number of tips per cell. n≥3 biological replicates. B. Imaris 9.9.0 and Image J software was utilized to quantify the average number of tips per cell. A box and violin plot illustrates significantly increased branching upon Pdlim5 or PalmD exogenous expression. Each dot represents a single neuron (n=10), and the levels of statistical significance, determined using one-way ANOVA, are indicated as P ≤ 0.0001 (****). n≥10 neurons. C. A bar graph compares negative-control GFP, exogenous PalmD expression, and knockdown of PalmD (siRNA-714; siRNA-716) based on Imaris 9.9 tool analysis (see Methods). The analysis reveals that PalmD acts as a modulator for branching function. The loss of PalmD is associated with a decrease in branching morphology, while exogenous expression of PalmD enhances branching. Each dot represents a single neuron (n=10), and the levels of statistical significance, determined using one-way ANOVA, are indicated as P ≤ 0.0001 (****), n≥10 neurons each from three biological replicates. D. Correlated total cell fluorescence was quantified using Image J software, indicating that knockdown of PalmD significantly reduces fluorescence intensity compared to the negative-control GFP or to positive-control PalmD-overexpressing neurons. n≥10 neurons in each of three biological replicates. Each dot represents a single neuron, and The violin plot quartiles represent data ranges and middle dark line represent median, and the statistical significance, determined using one-way ANOVA, is indicated as P ≤ 0.0001 (****). E. Sholl analysis was performed to analyze dendrite morphology in neurons under different conditions: GFP control, exogenous PalmD expression, and exogenous PalmD expression in the presence of knockdown of Pdlim5 (siRNA-992). Neurons overexpressing PalmD exhibit a significant increase in dendritic complexity compared to GFP-expressing cells. However, when Pdlim5 siRNA-992 is introduced alongside PalmD exogenous expression, there is a significant loss of branching function. n≥15 neurons in each of three biological replicates. The significance was assessed using a two-way ANOVA with Bonferroni post-hoc analysis. The levels of significance varied from P ≤ 0.0001 to P ≤ 0.05, depending on the radial distance from the soma., F. Representative binary images (see Methods) indicate the interdependency of Pdlim5’s and PalmD’s branching functions. In the absence of Pdlim5, PalmD no longer enhances branching morphology. Analogously, in the absence of PalmD, Pdlim5 displays a loss of branching phenotype. n=3 biological replicates. Scalebars 20µm.

## Notes

### Competing Interest Statement

The authors have declared no competing interest.

### Summary of Updates

We identified typographical errors in the manuscript(citations), and rectified them to improve the clarity for readers. We respectfully request permission to upload the revised manuscript version.

