## Supplementary figures and images for "“Role of a Pdlim5:PalmD complex in directing dendrite morphology”"

### Supplemental Data 1

Figure S3: PalmD, a novel partner of Pdlim5, associates with Pdlim5's LIM domain

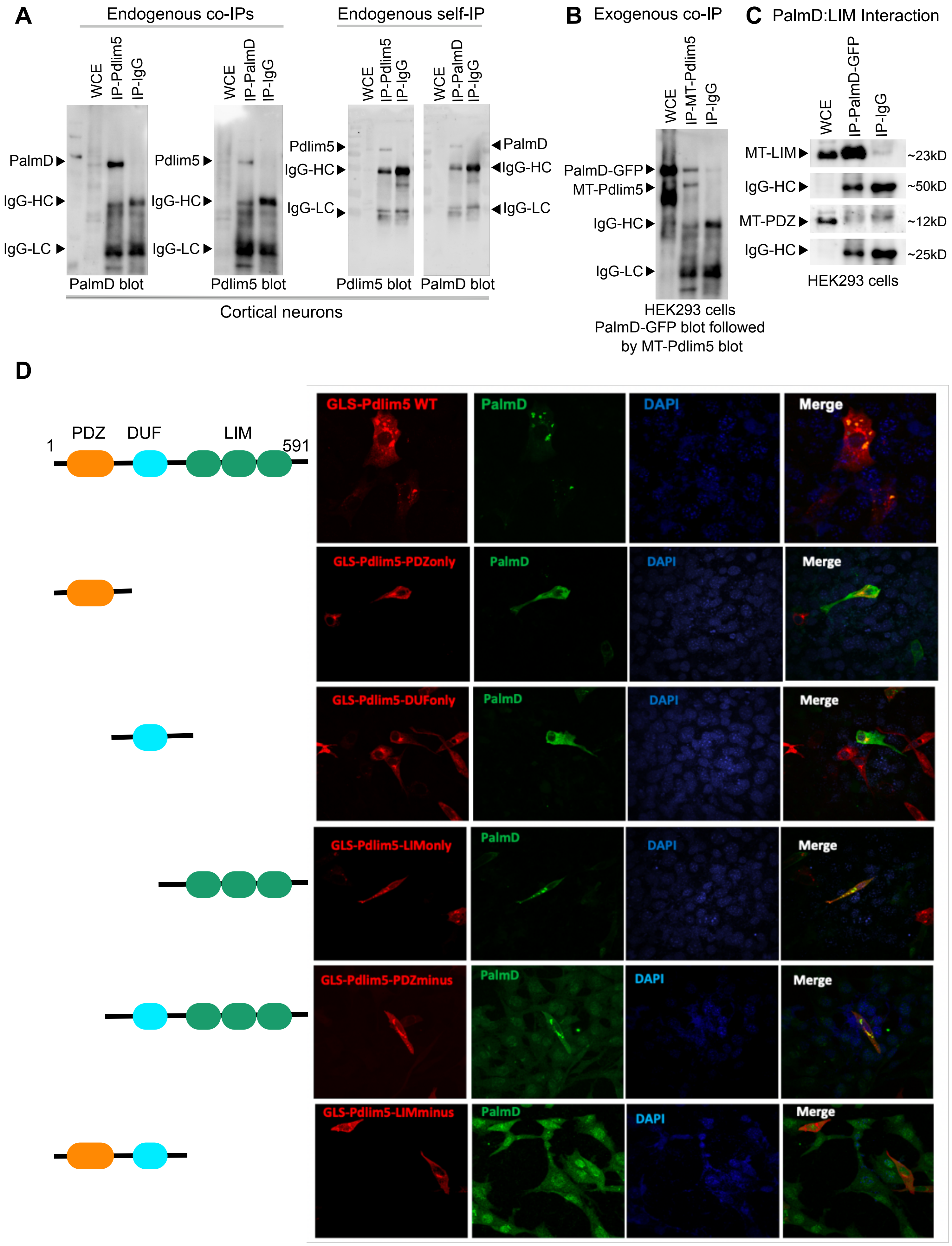

### Supplemental Figure 1. Pdlim5 conservation and function

Figure S1: Pdlim5 conservation and function

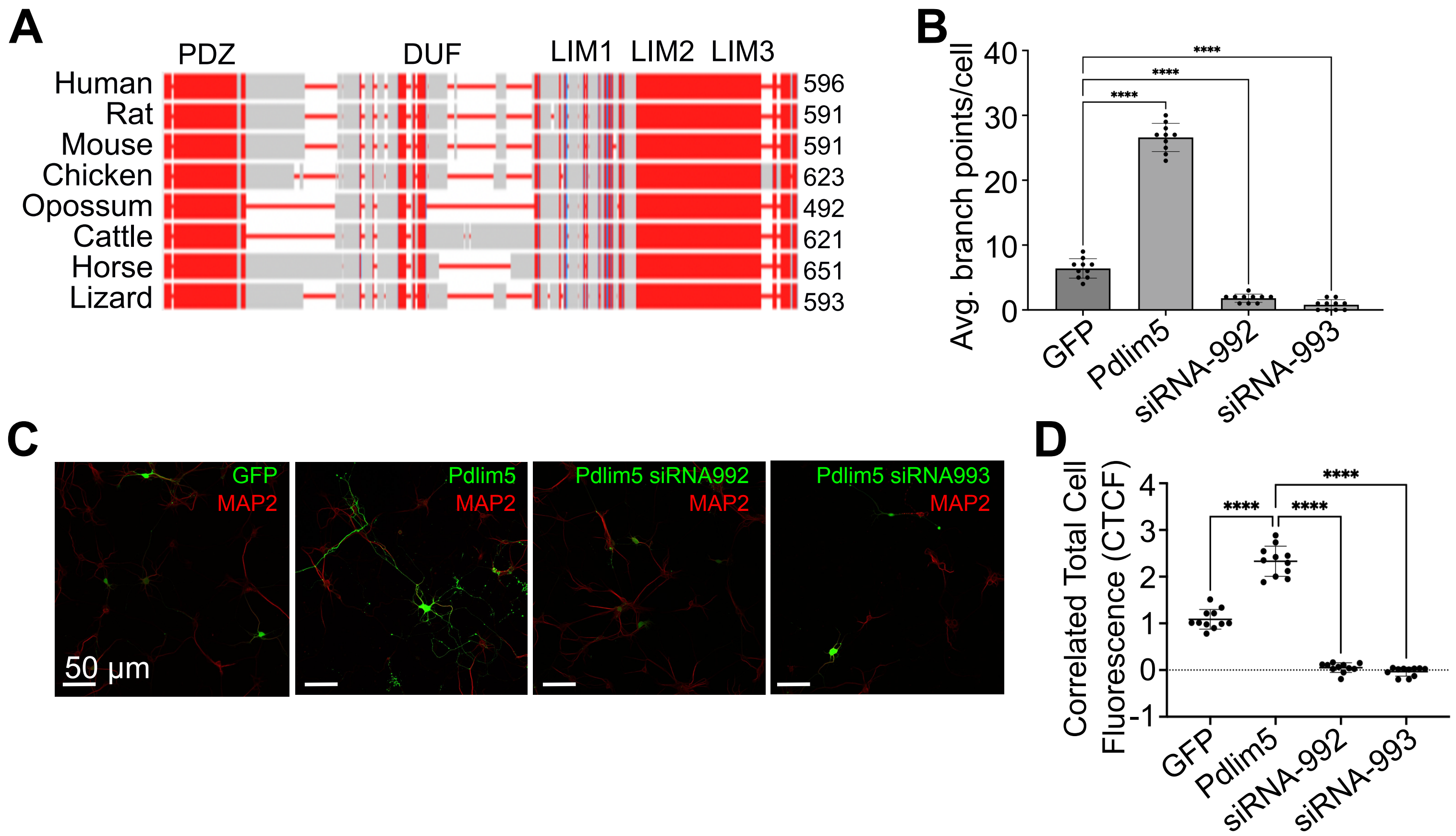

### Supplemental Figure 4. Function and functional dependency of Pdlim5: PalmD complex

#### Figure S4: Function and functional dependency of Pdlim5:PalmD complex

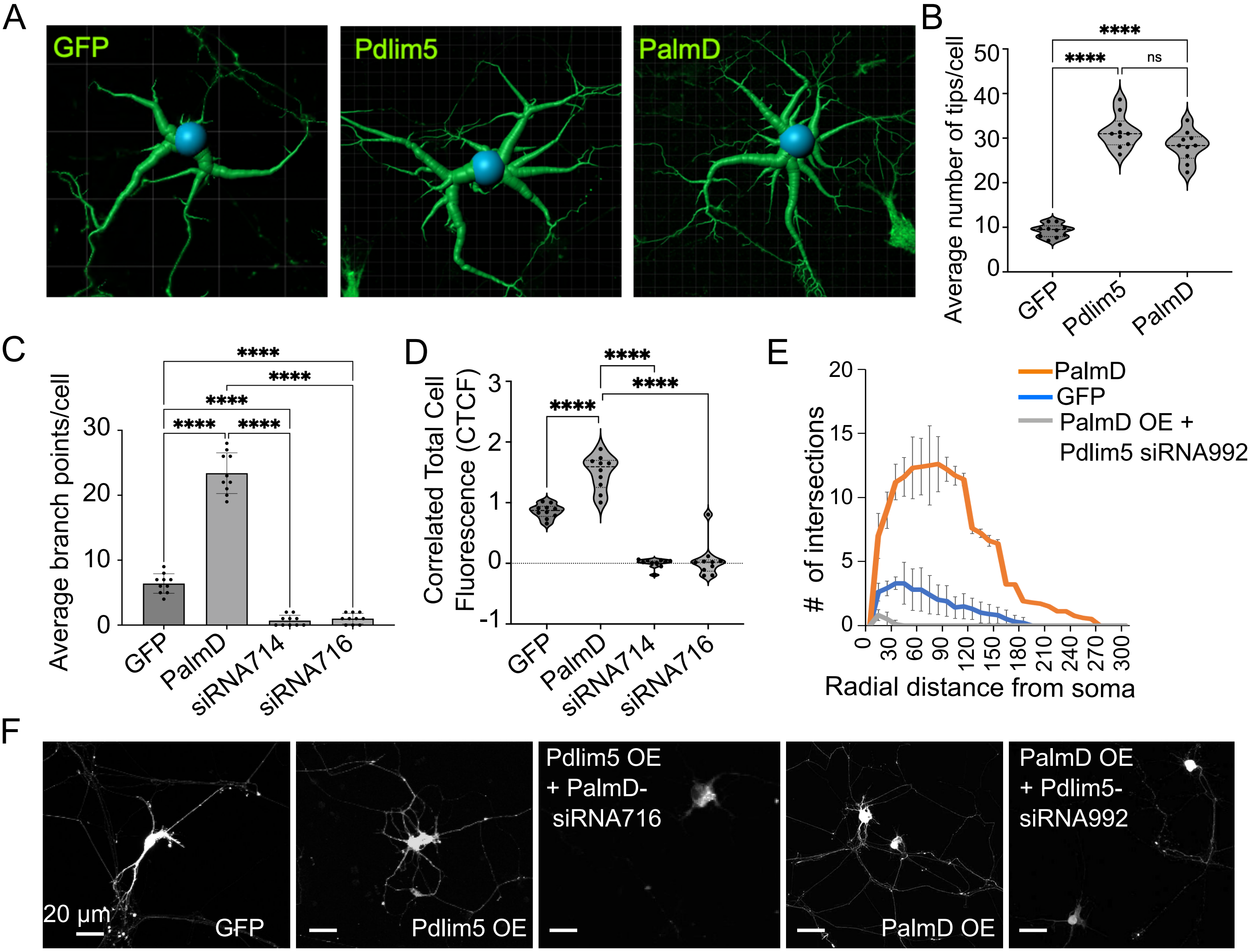
