## Supplemental Figure 2. Selected Pdlim5 associated candidates with known cytoskeletal functions for "“Role of a Pdlim5:PalmD complex in directing dendrite morphology”"

Figure S2: Selected Pdlim5 associated candidates with known cytoskeletal functions

A

Pdlim5 Y2H screen: Selected Prey (Gene Product) Findings and Roles

| | |
|---|---|
| PalmD | Linker of cytoskeleton to the plasma membrane. |
| Actn4 | alpha-Actinin4, cross-linker of actin microfilaments |
| Macf1 | Facilitates actin-MT interaction |
| Nin | Anchoring minus-end MTs. |
| Palld | Component of microfilaments. Cell shape control. |
| Pcnt | MT nucleation. Interacts with gamma-tubulin. |
| Sorbs2 | Potential link between Abl family kinases and actin. |
| Specc1l | Actin organization and MT stabilization. |
| Sptbn2 | Beta-III Spectrin. Actin binding cytoskeletal scaffold. |
| Sptbn4 | Beta-IV Spectrin. Actin binding cytoskeletal scaffold. |
| Vps18 | Vesicle trafficking endosome/ lysosome. |
